# Gut segments outweigh the diet in shaping the intestinal microbiome composition in grass carp *Ctenopharyngodon idellus*

**DOI:** 10.1101/356030

**Authors:** Wenwen Feng, Jing Zhang, Ivan Jakovlić, Fan Xiong, Shangong Wu, Hong Zou, Wenxiang Li, Ming Li, Guitang Wang

## Abstract

**ABSTRACT:** Although dynamics of the complex microbial ecosystem populating the gastrointestinal tract of animals has profound and multifaceted impacts on host’s metabolism and health, it remains unclear whether it is the intrinsic or extrinsic factors that play a more dominant role in mediating variations in the composition of intestinal microbiota. To address this, two strikingly different diets were studied: a high-protein, low-fiber formula feed (FF), and low-protein, high-fiber Sudan grass (SG). After a 16-week feeding trial on a herbivorous fish, grass carp, microbial profiles of midgut (M) and hindgut (H) segments of both groups were compared. Bacteroidetes were more abundant in the hindgut (T=-7.246, p<0.001), and Proteobacteria in the midgut (T=4.383, p<0.001). Fusobacteria were more abundant in the FF group (compared to the SG group, T=2.927, p<0.001). Bacterial composition was different (p<0.05) between the midguts of formula feed (M-FF) and Sudan grass (M-SG) groups, but not between the hindguts of two groups (H-FF and H-SG; p=0.269). PerMANOVA and VPA indicated that the gut segment contributed 19.8% (p<0.001) and 28% (p<0.001) of the variation of microbial communities, whereas diet contributed only 8.0% (p<0.001) and 14% (p<0.001), respectively. Overall, results suggest that intestinal compartments are a stronger determinant than diet in shaping the intestinal microbiota. Specifically, whereas diet has a strong impact on the microbiome composition in proximal gut compartments, this impact is much less pronounced distally, which is likely to be a reflection of a limited ability of some microbial taxa to thrive in the anoxic environment in distal segments.

**IMPORTANCE:** The impact of compositional dynamics of gut microbiota on host’s metabolism and health is so profound that the traditional idea of biological individual is increasingly replaced with "holobiont", comprising both the host and its microbiome. Composition of gut microbiota is strongly influenced by extrinsic (such as diet) and intrinsic (such as gut compartment) factors. Despite ample scientific attention both of these factors have received individually, their relative contributions in mediating the dynamics of the microbiome remain unknown. Given the importance of this issue, we set out to disentangle their individual contributions in a herbivorous fish, grass carp. We found that intestinal compartments are a stronger determinant than diet in shaping the intestinal microbiota. Whereas the impact of diet is strongly pronounced in proximal gut compartments, it appears that limited ability of some microbial taxa to thrive in the anoxic environment in distal segments strongly reduces the impact of diet distally.

---

Gastrointestinal tract of animals harbors an extremely diverse and complex microbial ecosystem (1–3). In the course of coevolution of gut microbiota and hosts, gut microbial community has become an integral component of the host (4, 5). Apart from contributing to the harvest of dietary nutrients that would otherwise be inaccessible to the host (6, 7) and to the education of the host’s immune system (8, 9), they also have profound impacts on host’s development and behavior (10, 11).

Although dominant members of gut microbiota are generally relatively constant (12), their overall composition is very variable, and strongly influenced by extrinsic and intrinsic factors (4, 13, 14), resulting in notable variability among individuals (3). Regarding the extrinsic factors, diet is known to be a major determinant of the microbial community composition in both terrestrial (15) and aquatic vertebrates (16). Among the intrinsic factors (e.g. gut physiology, host’s phylogeny or genotype), gut segments are a strong predictor of the composition of intestinal microbial communities in terrestrial mammals (4, 17, 18). However, it still remains unclear whether it is the host’s gut segment or dietary intake that plays a more dominant role in mediating variations in the composition of intestinal microbiota.

Characterization of the intestinal microbiota and their ecological function is relatively advanced in humans and model mammals (19, 20), but less well understood in fish (3). Intestinal microbiota of fish are believed to be less complex and less numerous than those of terrestrial vertebrates (9). Due to the tremendous importance of herbivorous grass carp (*Ctenopharyngodon idellus*) for the freshwater aquaculture and nearly global distribution (21), its intestinal microbiome has been studied extensively in recent years (3, 22, 23). Proteobacteria, Firmicutes, Bacteroides, Actinobacteria, and Fusobacteria are dominant in its intestine (23–25). Further investigations indicate that the intestinal microbiome of grass carp is likely to play an indispensable role in nutrient (especially polysaccharide) turnover and fermentation of the host (16, 26). Therefore, maintaining homeostasis of intestinal microbiota is likely to be essential for health and survival of grass carp..

The intestinal tract of grass carp is a simple coiled tube with eight convolutions, divided into three different segments according to its anatomical structure: foregut, midgut and hindgut (27). Theoretically, physiological functions should be distinct in different intestinal regions: foregut is believed to be responsible for the absorption of lipids and hindgut for pinocytotic uptake of macromolecules, including proteins (28, 29). However, most studies of intestinal microbiota in grass carp focused on the extrinsic factors, such as environment (3), geolocation (24), host’s diet (3, 30) and dietary supplementation application (31, 32), whereas studies of the roles of intrinsic factors, including the gut compartments, in shaping the intestinal microbiota in fish remain absent.

As diet is believed to be the most important force shaping the gut bacterial community in fish as well as other animals (33–36), we hypothesized that diet should outweigh the intestinal segments in shaping the composition of microbial populations in grass carp. To achieve this, we used two very different diets: formula feed (high-protein, low-fiber) and Sudan grass (high-fiber, low-protein), and sampled microbial populations of midguts and hindguts of both diet groups after the feeding experiment. Following this, we compared the microbial profiles of midgut and hindgut of both diet groups, and statistically tested the relative impacts of dietary intake and different gut segments on shaping the gut microbiota in the midgut and hindgut of grass carp. Therefore, the objectives of this work were two-fold: to infer differences in the microbial taxonomic composition among different intestinal compartments in grass carp, and to contribute to the understanding of relative contributions of diet and gut physiology on the microbial population structure in animals in general.

## RESULTS

### Bacterial community diversity

Community richness and diversity varied among gut segments and different diets (Table 1). All four richness and diversity indices were significantly higher in the midgut of both diet groups: M-FF (midgut-formula feed)>H-FF (hindgut-FF) (T_chao1_=4.954, p<0.01; T_ACE_=4.850, P<0.01; T_shannon_=4.938, P<0.01; Tsimpson 2. 326, p<0.05), and M-SG (midgut-Sudan grass)>H-SG (T_chao1_=3.393, p<0.01; T_ACE_=3.370, P<0.01; T_shannon_=5.379, P<0.01; T_simpson_=5.136, p<0.01). When considering each diet independently, community richness of the FF group was significantly higher than that of the SG group (T_chao1_=3.408, p<0.01; T_ACE_=3.582, p<0.01). Nevertheless, community diversity was not significantly different between FF and SG groups (T_shannon_=1.908, P=0.06; T_simpson_=0.841, p=0.403). The highest community richness and diversity indices were found in the M-FF group (Chao1=1351.30±345.69, ACE=1413.41±338.80, Shannon=6.66±2.03, and Simpson=0.91±0.16), while the lowest were found in the H-SG group (Chao1=527.37±413.70, ACE= 533.72±452.78, Shannon= 3.45±1.38, and Simpson=0.76±0.09).

**Table 1.** Summary of alpha diversity estimators for microbial communities of four groups. M-FF, midgut samples of the group fed on formula fed; M-SG, midgut samples of the group fed on Sudan grass; H-FF, hindgut samples of the group fed on formula fed; H-SG, hindgut samples of the group fed on Sudan grass.

| Group | Richness estimates |  | Diversity estimates |  | Good's coverage |
| --- | --- | --- | --- | --- | --- |
|  | Chao 1 | ACE | Shannon | Simpson | Mean±SD |
|  | Mean±SD |  | Mean±SD |  |  |
| M-FF | 1351.30±345.69 | 1413.41±338.80 | 6.66±2.03 | 0.91±0.16 | 0.96±0.01 |
| M-SG | 938.10±413.70 | 975.91±452.78 | 5.58±1.38 | 0.91±0.09 | 0.94±0.05 |
| H-FF | 796.19±345.69 | 850.99±338.80 | 4.03±2.03 | 0.81±0.16 | 0.98±0.01 |
| H-SG | 527.37±413.70 | 533.72±452.78 | 3.45±1.38 | 0.76±0.09 | 0.99±0.05 |

### Bacterial community composition

Using the diet+segment grouping, at the phylum level, Proteobacteria (46.63±19.7%), Firmicutes (23.52±19.47%), Fusobacteria (11.02±21.77%), Planctomycetes (7.70±8.70%), and Chloroflexi (3.28±3.34%) were dominant in the two midgut groups of samples (M-FF and M-SG; Figure 1). However, Bacteroidetes (29.79±24.22%), Proteobacteria (25.38%±21.40%), Firmicutes (21.52±12.76%), Fusobacteria (18.15%±21.29%) and Tenericutes (3.53±9.23%) were dominant in the two hindgut groups of samples (H-FF and H-SG; Figure 1). At the intestinal segment level, Bacteroidetes were significantly more abundant in the H group (T=-7.246, p<0.001), while Proteobacteria were more abundant in the M group (T=4.383, p<0.001). At the diet level, the dominant phyla in the FF group were Proteobacteria (33.56±19.02%), Fusobacteria (21.69±26.49%), Firmicutes (16.74±11.29%), Bacteroidetes (10.23± 17.77%), Planctomycetes (5.89±9.12%), and Tenericutes (4.83±9.38%), and dominant phyla in the SG group were Proteobacteria (38.46±26.54%), Firmicutes (28.30±18.67%), Bacteroidetes (20.10±25.71%), Fusobacteria (7.48±12.15%), Planctomycetes (2.17±3.71%), and Actinobacteria (1.12±1.45%). Statistical analysis indicated that Fusobacteria were significantly more abundant in the FF group than in the SG group (T=2.927, p<0.001). Bacteroidetes were more abundant in SG group than in FF group, but the difference was slightly above the selected statistical significance threshold (p=0.063).

**Figure 1.**
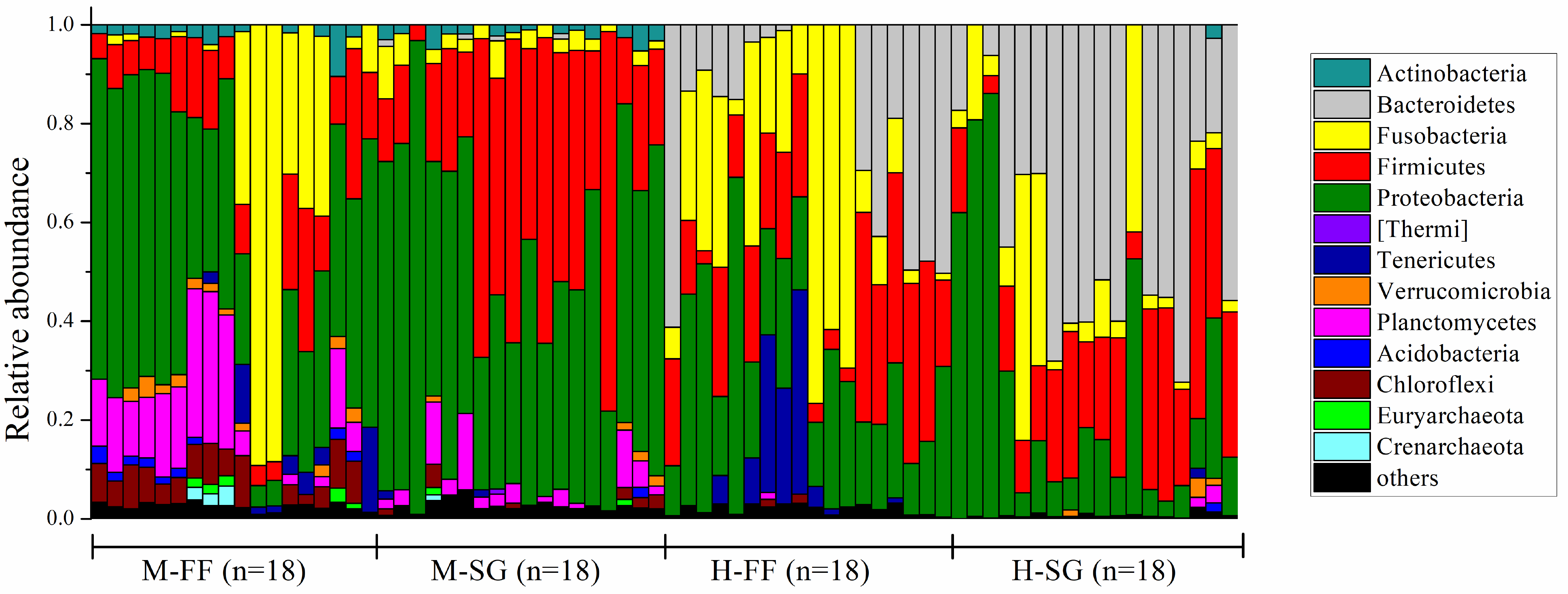
Composition of bacteria in four groups at the phylum level. Each bar represents the community of a sample. Only those phyla with mean relative abundance>1% are shown; whereas low abundance phyla were assigned to ’others’.

At the genus-level, the top ten most abundant genera differed among the four main sample groups (M and H, FF and SG; Table S1). On average, *Bacteroides* species were more abundant (P=0.076) in SG group (17.38±22.55%) than in FF group (9.05±16.17%). *Cetobacterium* were significantly higher (T=2.672, P<0.05) in FF group (18.53±25.83%) than in SG group (5.89±11.75%).

More than 700 bacterial taxa (genus or higher taxonomic level) significantly different (in terms of abundance) between the M-FF/H-FF and M-SG/H-SG group pairs were identified using Lefse with the LDA score value threshold set at 2.0 (Figures S1 and S2). In the FF group, Bacteroidetes (mostly Bacteroidia and *Bacteroides*), Erysipelotrichi and Aeromonadales (mostly Aeromonadaceae) were the most enriched taxa in the hindgut, whereas Desulfobacteria, Planctomycetes, and Pirelluales (mostly Pirelluaceae) were the most significantly enriched taxa in the midgut (Figure 2a). In the SG group, Bacteroidetes (mostly Bacteroidia and *Bacteroides*) and Aeromonadaceae were also the most enriched taxa in the hindgut, followed by Fusobacteriaceae, but Proteobacteria, Bacilli and Streptococcaceae (mostly *Streptococcus*) were the most significantly enriched taxa in the midgut (Figure 2b).

**Figure 2.**
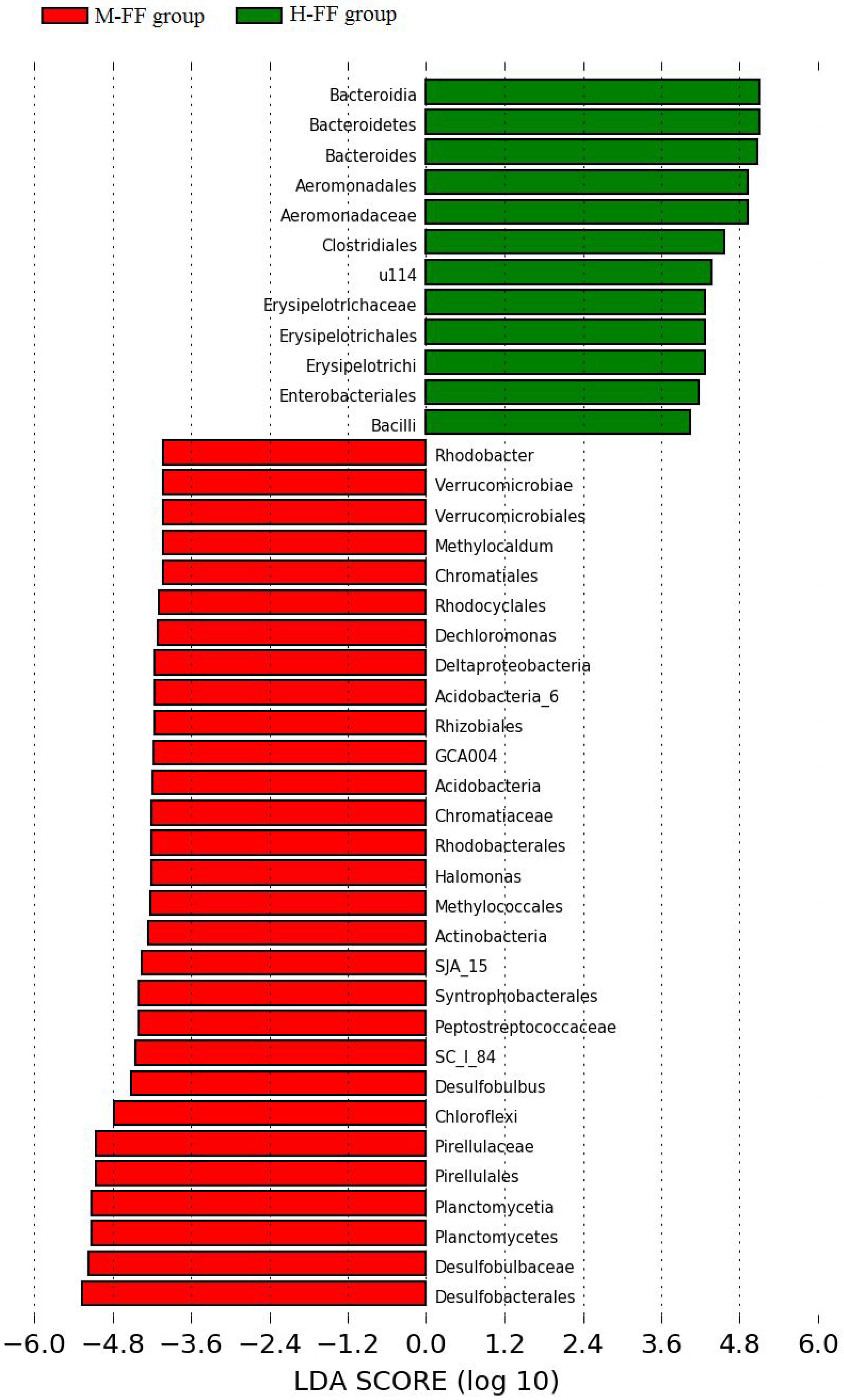
Bacterial taxa significantly different between the M-FF and H-FF groups (2a) or between the M-SG and H-SG groups (2b) identified by linear discriminant analysis coupled with effect size (LefSe) with LDA value set at 4.0.

### Relationships between bacterial communities of different gut segments and diets

A heatmap analysis at the family level showed that samples from the M group formed a single cluster, clearly distinct from the H group samples (Figure 3). PerMANOVA analysis revealed a significant difference (F=51.29, P=0.0001) in the composition of bacterial communities between M and H groups, but not between FF and SG groups (F=1.316, P=0.247). PerMANOVA with "adonis" algorithm indicated that grass carp gut segment contributed 19.8% (p<0.001) of the variation of gut bacterial communities, whereas diet contributed only 8.0% (p<0.001) (Table 2). Similarly, VPA analysis indicated that gut segments explain 28% (p<0.001) of the variation, and diet 14% (p<0.001).

**Figure 3.**
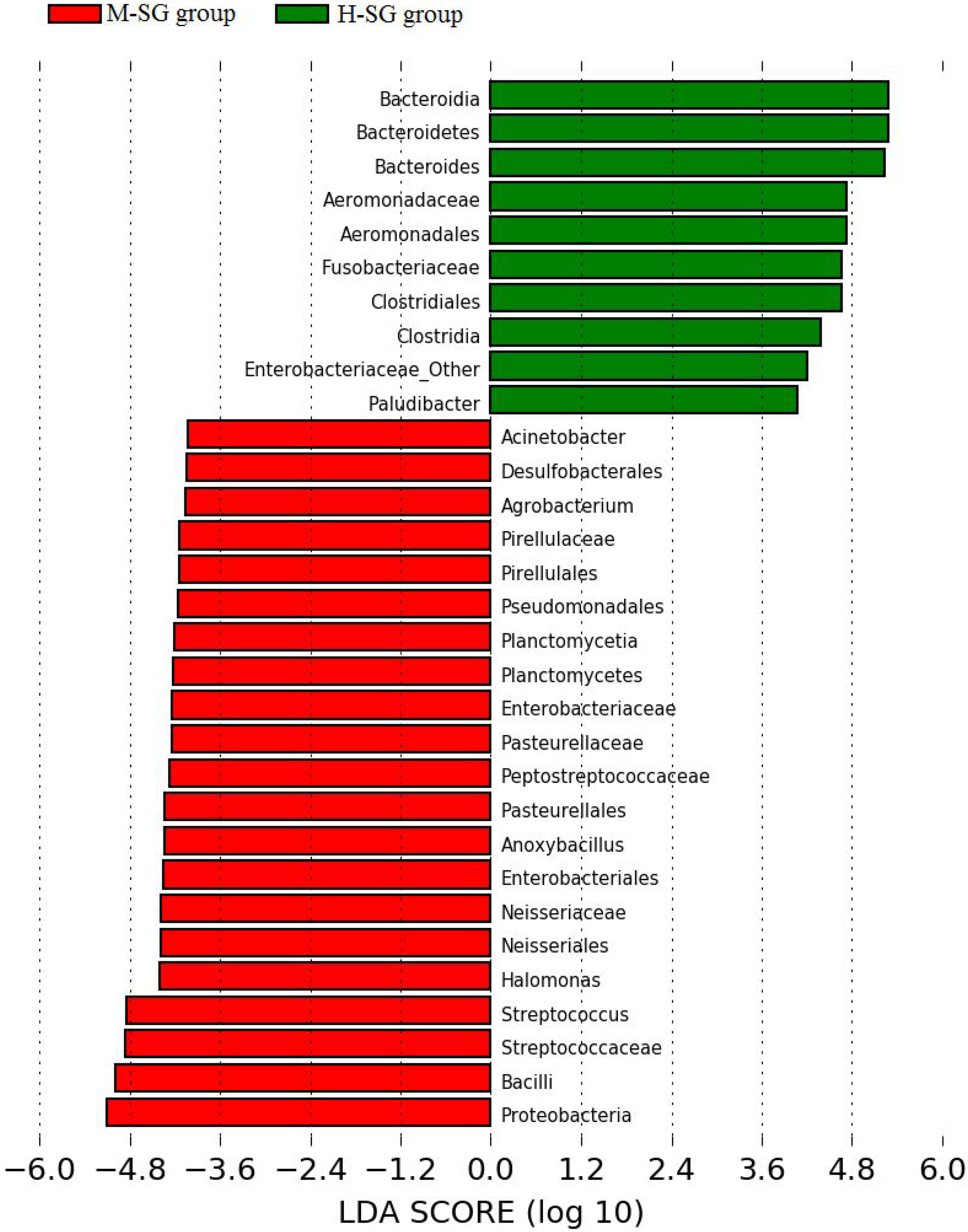
A heatmap plot depicting the relative percentage of each bacterial family with mean relative abundance>1% (variables clustering on the Y-axis) within each sample (X-axis clustering). Relationships among groups were determined using the Hierarchical clustering. In the heatmap, red colour indicates higher relative abundance, whereas blue colour indicates a lower relative abundance.

**Table 2.** Quantitative effects of gut segment and diet on the intestinal bacterial community assessed using permutational multivariate analyses of variance with Adonis function. R^2^ values represent the proportion of the community variation explained by each variable.

|  | Gut segment |  | Diet |  | Gut segment : Diet |  |
| --- | --- | --- | --- | --- | --- | --- |
|  | R<sup>2</sup> | p | R<sup>2</sup> | P | R<sup>2</sup> | P |
| Community variation | 0.198 | < 0.001 | 0.080 | < 0.001 | 0.041 | 0.001 |

PCoA results indicated that midgut and hindgut had significantly different bacterial compositions regardless of diet (p=0.0001 in all cases, PerMANOVA based on weighted Unifrac; Figure 4). After controlling for the gut compartment, we found a significant difference in bacterial composition between M-FF and M-SG samples (p=0.0324; Figure S3), but not between H-FF and H-SG samples (p=0.2688; Figure S4). We also determined the OTUs shared between these four groups of samples: M-FF and H-FF samples shared 1608 OTUs, M-SG and H-SG shared 1052, M-FF and M-SG shared 2401, and H-FF and H-SG groups shared 1272 OTUs (Figure S5).

**Figure 4.**
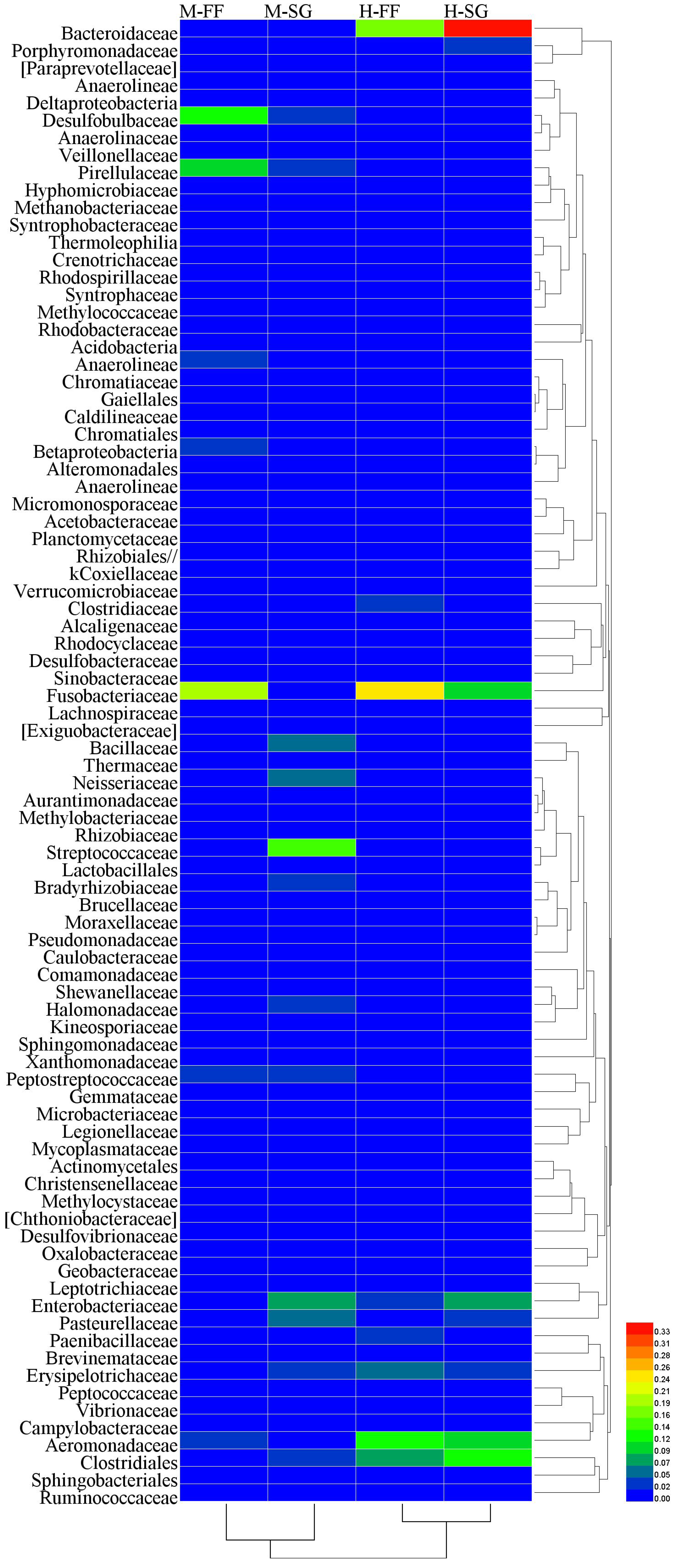
Principal coordinate analysis (PCoA) based on weighted UniFrac distances illustrating community dissimilarities over different gut segments and diet samples.

### Functional prediction of the midgut and hindgut microbiome

To infer the functional profiles of midgut and hindgut microbiomes, microbial 16S rRNA sequence data were analyzed by PICRUST to predict the dominant gene families. KEGG database level 2 query assigned the genes to 41 functional groups, predominantly to ’poorly characterized’, ’membrane transport’, and ’nucleotide metabolism’ (Figure 5). Nineteen gene families exhibited significant (p<0.05) differences between midgut and hindgut. The pathways these gene families were mainly associated with metabolic pathways: xenobiotics biodegradation and metabolism, nucleotide metabolism, metabolism of terpenoids and polyketides, metabolism of cofactors and vitamins, lipid metabolism, glycan biosynthesis and metabolism, energy metabolism, and carbohydrate metabolism. Some oxygen-independent pathways (especially fructose/mannose and starch/sucrose metabolisms) were also enriched in the hindgut samples (Figure S6).

**Figure 5.**
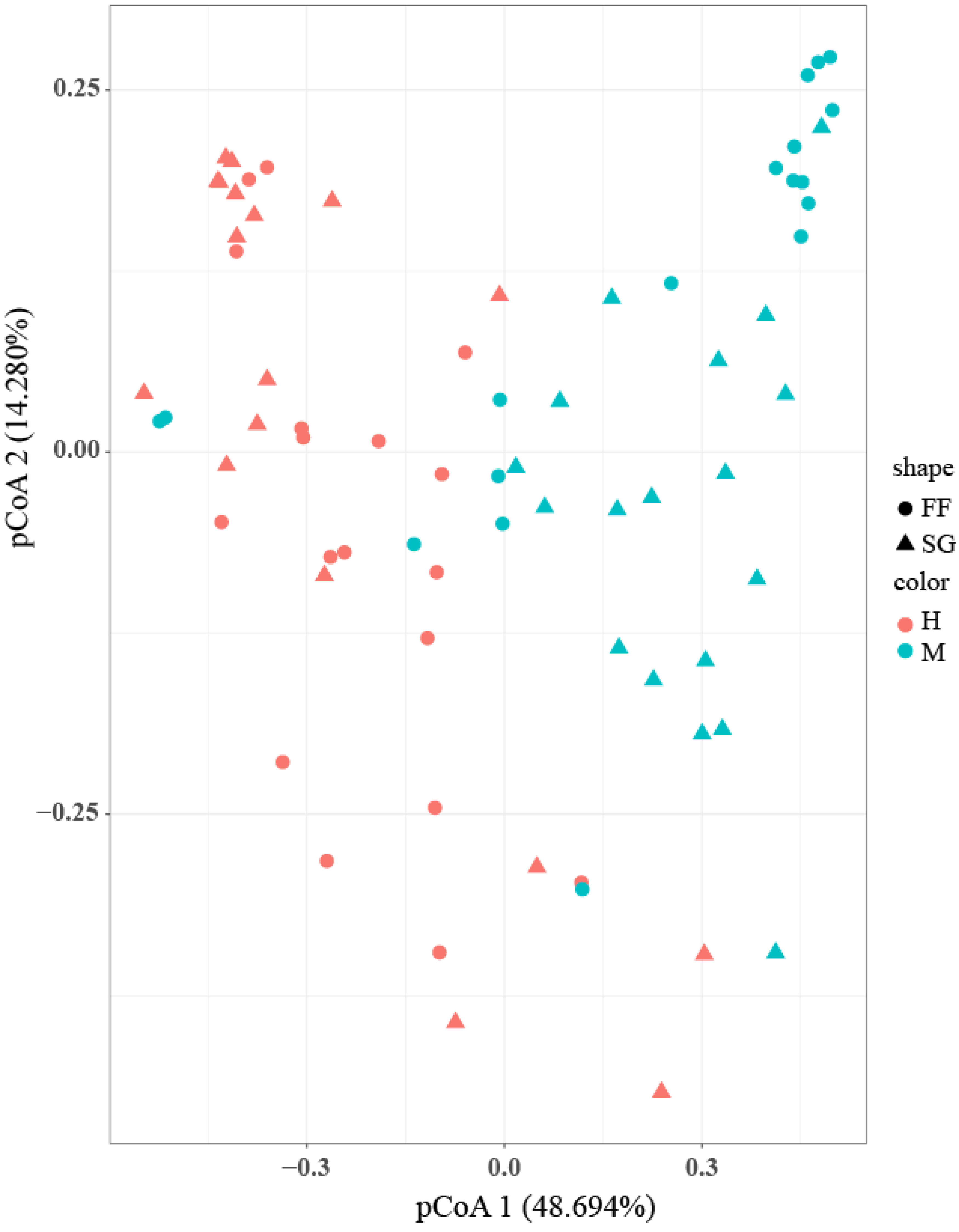
Functional profiling of midgut and hindgut microbial communities predicted by PICRUSt in the KEGG database (level 2). The significance level is indicated by *p<0.05; **p<0.01; ***p<0.001.

## DISCUSSION

Substantial research has been carried out in recent decades to better understand the complexity and diversity of gut microbiota in fish (22, 23, 37). Diet is known to be a very important factor influencing the intestinal bacterial composition. For example, in the Atlantic cod (*Gadus morhua* L), gram-positive *Brochothrix* and *Carnobacterium* were dominant in the gut of a fishmeal diet-fed fish, *Psychrobacter* dominated in the bioprocessed soy bean meal group, and *Carnobacterium*, *Chryseobacterium* and *Psychrobacter glacincola* dominated in the soy bean meal diet group (35). However, the impact of different gut compartments on the bacterial composition remains unstudied in fish.

Our study provides a detailed comparison of bacterial communities in different gut segments in a herbivorous fish, in combination with two strikingly different diets. Heatmap analysis indicated that midgut samples from both diet groups formed a single cluster, significantly different from the hindgut samples of both diet groups. This suggests that the composition of microbiome was impacted more substantially by the gut compartment than by the diet. However, large SD values observed in all of these analyses, as well as comparison with previous studies of this species (16), indicate that individual variability also plays a major role in determining the microbial composition.

This dramatic difference in the microbiome composition between midgut and hindgut may be related to gut morphology and physicochemical conditions (38, 39). Obligate anaerobes, including *Bacteroides* (Bacteroidetes), Fusobacteriaceae (Fusobacteria), and Clostridiales and Erysipelotrichaceae (Firmicutes), were significantly more abundant in hindgut samples than in midgut samples. Proteobacteria, however, were more abundant in the midgut samples. Metagenomes also revealed increasing prevalence of anaerobic metabolism in hindgut in comparison to midgut, which included fructose and mannose metabolism, galactose metabolism, and starch and sucrose metabolism. The observed shift towards obligate anaerobes is expected, as the hindgut is characterized by extremely low oxygen concentrations in most animals (40). *Bacteroides* was also reported as the most abundant taxon in the distal gut segments of a broad spectrum of animal species, from mammals (sheep rectum) (18) to insects (*Pachnoda ephippiata*, distal gut) (41). However, dominant taxa varied among the proximal gut samples of these three species: *Streptococcus* in sheep jejunum (18), aerobic Actinobacteria in the midgut of *P. ephippiata* (41), and Proteobacteria in grass carp. Therefore, oxygen levels are the most likely explanation for the observed significant difference in the bacterial composition between bacterial communities of midgut but not hindgut samples of the two diet groups: in an aerobic environment, diet is the major factor determining the microbial composition, but as the environment turns anaerobic, it becomes hospitable only for a limited number of microbial taxa, resulting in shrinking microbial richness and diversity indices.

As diet is believed to be the most important force shaping the gut bacterial community (34–36), we also studied the impacts of two very different diets: Sudan grass and formula feed. When each diet was considered independently, bacterial community richness of the FF group was significantly higher than that of the SG group. Bacteroidetes (non-significantly) and *Bacteroides* were more abundant in the SG group. The genome of *Bacteroides* is enriched in glycoside hydrolase and polysaccharide lyase genes, targeting the degradation of the plant cell wall polysaccharides (16). Hence, high abundance of *Bacteroides* in the SG group probably reflects the high proportion of fiber in this diet. Similarly, gut microbiomes of high-fiber diet consuming humans are highly enriched in Bacteroidetes (42). On the other hand, the *Cetobacterium* genus was significantly more abundant in the FF group (compared to SG group). This genus is known to be in a positive correlation with the production of acetic and propionic acids through peptone and glucose fermentation (43), and numerous gene families associated with protein digestion (peptidases) are present in the genome of *C. somerae*, which is an indigenous bacterium in the digestive tract of freshwater fish (16). This could be an explanation behind the high abundance of this microbe in high-protein formula feed diet-fed fish (16, 25).

## Conclusions

Composition of the intestinal bacterial community is determined by a large number of factors, including the host’s diet, gut compartment, life history, genetics, and environmental factors (3, 4), but diet is believed to outweigh the host’s genotype in shaping the gut microbiota (33). We found that the opposite is true for gut segments: both PerMANOVA and VPA analyses indicated that gut segments explain a higher proportion of the variation in intestinal microbiota than the diet. Despite the large individual variability observed, these results indicate that we can reject our working hypothesis, as intestinal anatomy and physiology appear to be a stronger determinant in shaping the intestinal microbiota than host’s diet. Apart from the understanding of bacterial functions in different gut segments, this finding also bears relevance for the interpretation of past studies and design of future studies of intestinal microbiota, which should pay close attention to the intestinal segment variability.

## MATERIALS AND METHODS

### Sample collection

Juvenile fish were purchased commercially and kept in artificial earthen ponds in Huanggang City, Hubei Province, China, from April to August, 2015. Six ponds (with 30 fish in each pond; 1.5-2.0 m depth, 100m^2^ surface) were divided into two groups: one group was fed the Sudan grass diet (SG group) and the other was fed the formula feed diet (FF group). The Sudan grass diet contained 29% crude fiber and 10.37% crude protein, whereas the formula feed diet contained 6.9% crude fiber and 40.45% crude protein (44). The fish were fed to apparent satiation twice a day (8:00 and 16:00 o’clock). After the feeding experiment (16 weeks), six grass carp specimens were randomly collected from each pond (6×6=36 specimens). Fishes were euthanized in buffered MS-222 at 250 mg/L concentration, measured (weight and length) and immediately dissected in sterile conditions. Body length was 30.67±2.73 cm and weight was 486.57±126.99g. Intestines were divided into segments as described before (27), the entire content of midgut and hindgut collected, separately placed into labelled 25mL polypropylene centrifuge tubes, frozen provisionally in a portable refrigerator, transported to laboratory within six hours and stored at -80 °C. This study has been reviewed and approved by the ethics committee of the Institute of Hydrobiology, Chinese Academy of Sciences.

### DNA extraction, PCR amplification and sequencing

Genomic DNA was extracted from 72 samples (36 specimens× 2 gut segments) using QIAamp DNA stool mini kit (Qiagen, Germany) according to the manufacturer’s instructions. DNA concentrations were estimated using a Nanodrop 8000 Spectrophotometer (Thermos, USA). Obtained DNA samples were used for the amplification of bacterial V4-V5 16S rRNA gene region with universal barcode primers 515F (5’-GTGYCAGCMGCCGCGGTA-3’) and 909R (5’-CCCCGYCAATTCMTTTRAGT-3’) (45). PCR reaction mix (25μL) contained 0.5U of the Phusion high-fidelity DNA polymerase (New England Biolabs, Beijing China Ltd), 5×Phusion GC buffer, 5mM dNTP, 20μM primers and 50ng DNA. An initial denaturation at 98°C for 30s was followed by 25 cycles (98°C for 10s, 55°C for 20s and 72°C for 20s) and the final extension step for 10min at 72°C.PCR products were purified using AidQuick Gel Extraction Kit (Aidlab Biotech, Beijing, China). Purified samples were sequenced using Novogene bioinformatics technology on the Illumina Hiseq 2500 platform.

### Bioinformatic and statistical analyses

Raw sequenced data were analyzed using QIIME Pipeline-version 1.7.0 (46). Each sample was distinguished according to its unique barcode sequence (barcode mismatches=0). The first processing step was merging paired-end reads using FLASH-1.2.8 program (47). Only the merged sequences with high-quality reads (length>300 bp, without ambiguous base N, and average base quality score>30) were used for further analyses. Sequence chimeras were removed using the UCHIME algorithm (48). All sequences were grouped as operational taxonomic units (OTUs), applying a 97% identity threshold. Singletons and chloroplasts were filtered out. The sequence number of each sample was normalized to 11000 sequences. All sequences analyzed in this study can be accessed in the SRA database under the accession number SRP 131857.

Samples (n=72) were grouped using different criteria, diets (FF + SG; n=36), gut segments (Midgut + Hindgut, n=36), diet+segment (H-FF, M-FF, H-SG; M-SG; n=18), and statistically analysed. Alpha diversity indices of gut bacterial communities, including community richness (Chao1 and Ace) and diversity (Shannon and Simpson), were calculated using the QIIME package. To evaluate the beta diversity and visualize differences in the bacterial community structure, principal coordinates analysis (PCoA) was conducted using the weighted UniFrac distance. To identify relative abundance of bacterial biomarker taxa at the genus level between the midgut and hindgut of different diet groups, linear discriminant analysis coupled with effect size (Lefse) was employed on the Huttenhower laboratory Galaxy website (http://huttenhower.sph.harvard.edu/galaxy/) (49). Default logarithmic (LDA) score value thresholds were set at 2.0 (to identify all significantly different taxa) and 4.0 (to generate publishable figures focusing only on the most significantly different taxa). Venn diagram was used to display shared OTUs between different parts of the intestine and different diets (50). To reveal the similarities and differences among groups, a heatmap plot was constructed on the basis of the mean relative abundance of bacterial families which exceeded 0.1% in each sample. PICRUST1.0 (51) and KEGG database were used to explore functional profiles of the bacteriome in different gut segments. Bar graph was constructed using OriginPro 8.5 (52), and STAMPv2.1.3 (53) was used for statistical analyses of functional profiles. Statistical differences were calculated using Welch’s t-test with Bonferroni correction, with statistical significance threshold set at 0.05. Permutational multivariate analyses of variance (PerMANOVA) were performed using PAST 2.16 (54) to assess the significance of differences in the bacterial community structure among different groups, based on weighted UniFrac distance. PerMANOVA with "adonis" procedure was used to evaluate whether the diet and the gut segment significantly affected the bacterial community structure of grass carp. Variance Partitioning Analysis (VPA) was used to evaluate the contribution of gut segments and diets to the microbial community variance.

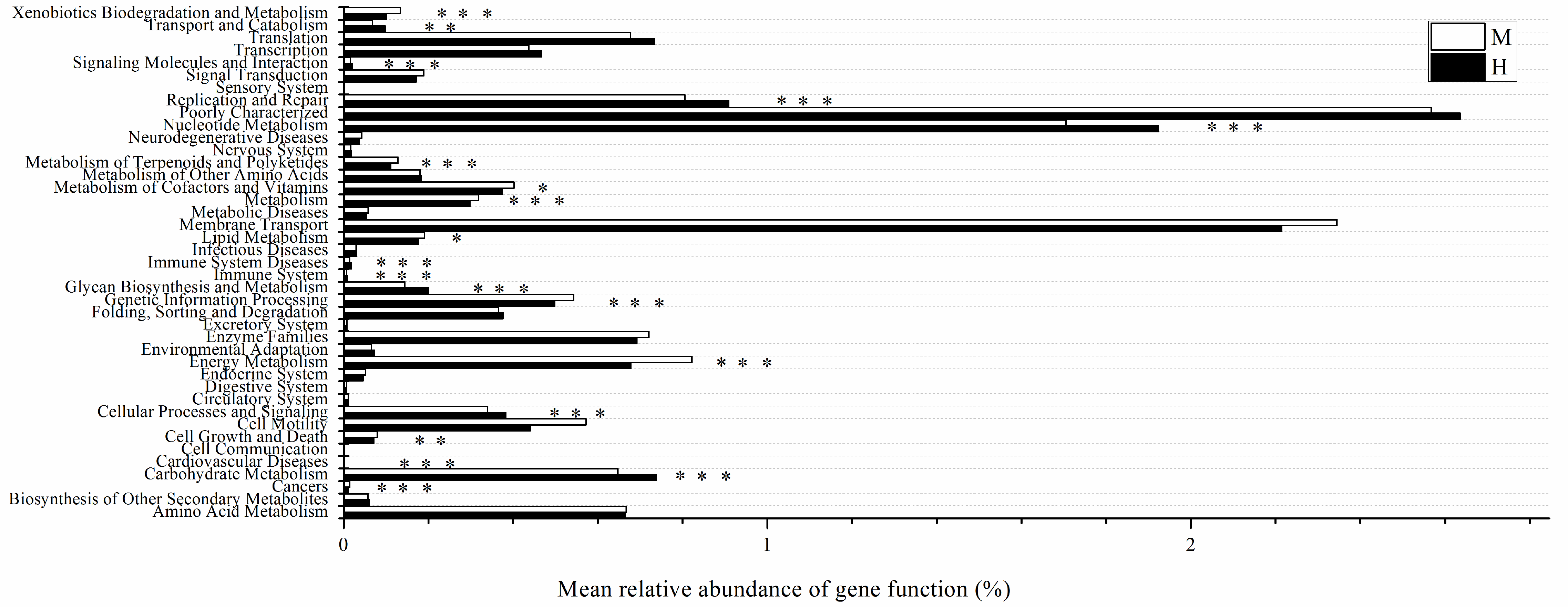

## ACKNOWLEDGMENTS

This work was supported by the National Natural Science Foundation of China (grants No. 31272706 and No. 31372571); and the earmarked fund for the China Agriculture Research System (No. CARS-45-08).

## DATA ACCESSIBILITY

All sequences analyzed in this study can be accessed in the SRA database under the accession number SRP 131857 (https://www.ncbi.nlm.nih.gov/Traces/study/?acc=SRP131857).

## AUTHOR CONTRIBUTIONS

The author contributions are as follows: S.G.W. and G.T.W. were principal investigators and contributed to the study design, acquisition of funding and overseeing the study, interpretation of data and manuscript editing. W.W.F. and J.Z. were in charge of the design, data collection, analysis and interpretation of data and manuscript drafting. F.X. contributed to data analysis and interpretation. I.J. was in charge of quality control, interpretation of the data and co-wrote the manuscript. H.Z., W.X.L. and M.L. were in charge of study coordination, quality control, and manuscript editing.

